# A genome-wide case-only test for the detection of digenic inheritance in human exomes

**DOI:** 10.1101/2020.02.06.936922

**Authors:** Gaspard Kerner, Matthieu Bouaziz, Aurélie Cobat, Benedetta Bigio, Andrew T Timberlake, Jacinta Bustamante, Richard P Lifton, Jean-Laurent Casanova, Laurent Abel

## Abstract

Whole-exome sequencing (WES) has facilitated the discovery of genetic lesions underlying monogenic disorders. Incomplete penetrance and variable expressivity suggest a contribution of additional genetic lesions to clinical manifestations and outcome. Some monogenic disorders may therefore actually be digenic. However, only a few digenic disorders have been reported, all discovered by candidate gene approaches applied to at least one locus. We propose here a novel two-locus genome-wide test for detecting digenic inheritance in WES data. This approach uses the gene as the unit of analysis and tests all pairs of genes to detect pairwise gene x gene interactions underlying disease. It is a case-only method, which has several advantages over classic case-control tests, in particular by avoiding recruitment and bias of controls. Our simulation studies based on real WES data identified two major sources of type I error inflation in this case-only test: linkage disequilibrium and population stratification. Both were corrected by specific procedures. Moreover, our case-only approach is more powerful than the corresponding case-control test for detecting digenic interactions in various population stratification scenarios. Finally, we validated our unbiased, genome-wide approach by successfully identifying a previously reported digenic lesion in patients with craniosynostosis. Our case-only test is a powerful and timely tool for detecting digenic inheritance in WES data from patients.

**Significance statement:** Despite a growing number of reports of rare disorders not fully explained by monogenic lesions, digenic inheritance has been reported for only 54 diseases to date. The very few existing methods for detecting gene x gene interactions from next-generation sequencing data were generally studied in rare-variant association studies with limited simulation analyses for short genomic regions, under a case-control design. We describe the first case-only approach designed specifically to search for digenic inheritance, which avoids recruitment and bias related to controls. We show, through both extensive simulation studies on real WES datasets and application to a real example of craniosynostosis, that our method is robust and powerful for the genome-wide identification of digenic lesions.

## INTRODUCTION

Next-generation sequencing (NGS) is now widely used and is gradually being optimized for the detection of rare and common genetic variants underlying human diseases (1–3). These advances, including whole-exome sequencing (WES), in particular, have led to major new findings in the field of human genetics, particularly for rare and common monogenic disorders (4–12). The growing number of reports of incomplete penetrance or variable expressivity of monogenic disorders suggests that additional genetic contributions, other than the mono- or biallelic causal lesions, may contribute to clinical manifestations and outcome (13, 14). Digenic inheritance (DI) is the simplest genetic model of this type with alleles at two different loci being necessary and sufficient to determine disease status (15, 16). The recently established Digenic Diseases DAtabase (DIDA) contains detailed information about DI for 258 reported digenic combinations, corresponding to 54 conditions, since 1994 (17). Well known examples relate to genetic modifier (GM) variants influencing the expression of the clinical phenotype caused by a primary disease-causing mutation. Cystic fibrosis (CF) is a classic example of a monogenic disease for which several GM variants have been identified. An elegant WES-based study showed that two low-frequency (minor allele frequency [MAF] < 5%) missense variants of *DCTN4* were associated with the severity of pulmonary *Pseudomonas aeruginosa* infections in CF patients (18). One remarkable example of DI explaining incomplete penetrance was recently provided for craniosynostosis. Timberlake et al. (2016) found a highly significant enrichment in rare damaging *SMAD6* mutations in patients with craniosynostosis (n=191). However, variants were also carried by 13 asymptomatic family members. The authors thus showed that a common variant close to *BMP2*, a *SMAD6*-related gene, accounted for almost all the observed incomplete penetrance.

Only 1% of the 5,442 traits listed in OMIM as single-gene disorders are also known to display DI and are listed in DIDA. Interestingly, all the lesions known to be caused by defects with DI were discovered in candidate gene studies, rather than through unbiased GW statistical tests. In some cases, as for the cystic fibrosis example cited above, the defects were identified by single-gene analyses of patients with known disease-causing variants at the primary causal locus (18). The craniosynostosis example is unique in that its discovery involved a combination of GW single-gene analysis with prior knowledge of a common variant from genome-wide association study (GWAS) data (19, 20). For genetically heterogeneous diseases, such as Alport syndrome, for which there are three known disease-causing genes, long-QT syndrome and Bardet-Biedl syndrome, each with more than a dozen disease-causing genes, the proven digenic combinations display various modes of dominance and involve the known disease-causing genes (21, 22). However, other GM genes may be hidden among genes with an unknown functional impact on disease, or even genes with no detectable main effect. Similarly, many heritable conditions masked in apparently sporadic cases, for which the genetic etiology remains unknown, may be due to DI.

There is, therefore, a need for two-locus GW methods for the detection of DI in NGS data. WES is a NGS technique focusing on sequencing of protein-coding exons. It is currently the most cost-effective NGS technology, as variants with a strong effect are more likely to affect protein-coding sequences than non-coding sequences (23–25). Very few methods have been developed for detecting gene x gene interactions in the general context of rare variant association studies; all techniques to date are based on case-control designs (26–28). Here, we propose a case-only approach to specific searches for DI. This design avoids the need for control recruitment and the associated bias. Furthermore, case-only approaches have been shown to be more powerful than classic case-control tests when common variants are tested for interaction, particularly in the context of GWAS (29–33). Our novel approach is based on the aggregation of rare variants within a gene as the unit of analysis, overcoming the lack of power inherent to studies of rare variants. It also greatly decreases the computer time required for interaction analyses, by testing pairwise combinations at the gene level.

## MATERIAL AND METHODS

### The variant aggregation model

A strategy commonly used for low-frequency variants from NGS data involves tests based on the aggregation of variants within a genomic region. Several types of tests are used for this purpose: burden tests, adaptive burden tests, variance-component tests and combinations of these three classes (34). Here, we propose a method based on the classic collapsing of variants within the unit of a gene. This approach optimizes statistical power under a hypothesis of genetic homogeneity, whilst making it possible to assess actual gene x gene interactions with a number of tests corresponding to the number of possible two-way combinations of genes. In this study, the aggregation of variants within a gene is based on the methodology of a class of burden tests known as the “cohort allelic sums test” (CAST). Formally, for each gene *j* and a given subset of variants *S*_*j*_ observed within this gene, if *n* is the number of individuals studied, we consider the following vector (*g*_*j*1_, …, *g*_*jn*_) denoted *G*_*j*_. For each *i* = 1, …, *n, g*_*ji*_ is then defined as follows:

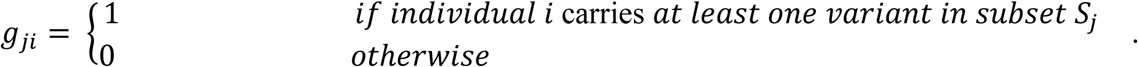

The term “carries” depends here on the biological inheritance model. For example, in a dominant model, *g*_*ji*_ = 1 if individual *i* harbors at least one copy of at least one variant allele from the set of variants studied *S*_*j*_ within gene *j*. In addition, the choice of *S*_*j*_ may be based on different features at the variant level, such as the MAF or functional impact prediction, as described below.

### The case-control design for interaction

Using this notation, data for genes *k* and *j* in a case-control dataset, with a binary disease status D, can be summarized into two 2×2 contingency tables, one for affected individuals (cases, D=1) and one for unaffected individuals (controls, D=0), as in Table 1. Based on these tables, let 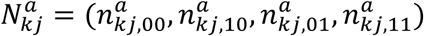 be a vector of the observed numbers of carriers for gene *k* and gene *j* among cases, such that, for example, 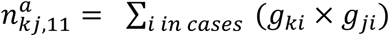. Similarly, we define 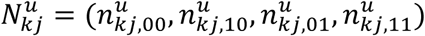 as a vector of the observed numbers of carriers for gene *k* and gene *j* among controls. The odds ratios for cases and controls, respectively, for genes *k* and, are defined as follows:

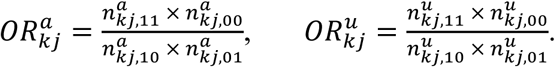

**Table 1.** Contingency table of carriers of rare variants for a given pair of genes *k* and *j* for affected and unaffected individuals.

| | Gene $k$ | |
| --- | --- | --- |
|  | Carriers | Non carriers |
| Gene $j$ | | |
| Carriers | $n_{kj,11}^i$ | $n_{kj,01}^i$ |
| Non carriers | $n_{kj,10}^i$ | $n_{kj,00}^i$ |

<sup>a</sup> $i = \{a, u\}$ . When $i = a$ , $n$ stands for the number of affected individuals, when $i = u$ , $n$ stands for the number of unaffected individuals.

Classic statistical analyses of interaction are based on the comparison of 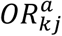 and 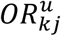. More specifically, the following classic case-control logistic regression model is often used to test for interaction:

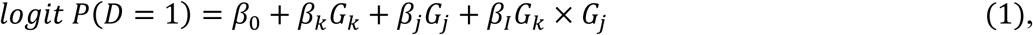

where it can be shown that the interaction coefficient, *β*_*I*_ equals 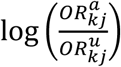. This model also takes main effects into account, by considering coefficient terms for each gene (*β*_*k*_ and *β*_*j*_). In addition, specific covariates, such as principal components (PCs), can easily be introduced into the model. Including a matrix of covariates *X* and a vector *C* of coefficients, the full logistic regression model takes the following form:

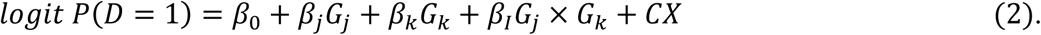

Subsequently, the null hypothesis of no interaction *β*_*I*_ = 0 can be tested in a likelihood ratio test (LRT) with one degree of freedom, in the presence or absence of main genetic effects and/or covariate effects.

### The case-only model

Interactions can also be assessed by focusing exclusively on cases, such that all the information is provided by the 2×2 contingency table for affected individuals (Table 1). In this situation, the standard full logistic regression model to test for interaction between genes G_k_ and G_j_ is now written as

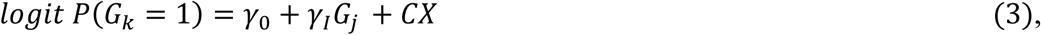

where *γ*_*I*_ is equal to 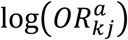, *X* is a matrix of covariates and *C* a vector of coefficients. As before, a LRT can be used to test the null hypothesis *γ*_*I*_ = 0.

Under the assumption that vectors G_k_ and G_j_ are not correlated, implying, in particular, that variants of the two genes are not in linkage disequilibrium (LD), a deviation from 1 of 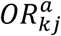 indicates interaction. In addition, if the disease is rare, 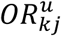 is close to 1, and, consequently, *β*_*I*_ is approximately *γ*_*I*_. The advantages of this test over case-control tests have been extensively studied theoretically (29, 33), in particular the gain of power. This gain stands from the nature of the estimators of the interaction coefficients of both designs. These estimators depend either on the ratio 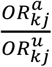 for the case-control or only on 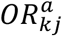 for the case-only test. The asymptotic variances of the estimators are the sum of the reciprocal counts of Table 1, either for both affected and unaffected subjects (case-control design), or for affected individuals only (case-only) (29). Hence, the variance of the estimator of the case-control interaction coefficient has a larger variance leading to a less powerful test. The advantages include also the absence of a need to recruit controls, which, in addition to saving time and reducing costs, avoids the problem of the misclassification of individuals with the unaffected phenotype. The only known limitation of this test is that it assumes independence in the general population of the variants tested. In fact, our type I error analyses revealed possible sources of violation of this assumption in the context of WES data that, to our knowledge, had never before been considered.

### Samples

For the simulation study we worked on real exome data, using samples from the 1000 Genome project (1000G) populations, and a subset of our in-house exome database, the Human Genetics of Infectious Diseases (HGID) database. Six populations from the 1000G database were used: four European populations — the Iberian population in Spain (IBS, *n*=107), Toscani in Italy (TSI, *n*=107), British in England and Scotland (GBR, *n*=91) and Finnish in Finland (FIN, *n*=99) — and two Asian populations of Chinese origin —Southern Han Chinese (CHS, *n*=105) and Chinese Dai in Xishuangbanna, China (CDX, *n*=93). From the HGID database, which includes data for > 4,000 individuals of various ethnic origins, including patients suffering from severe infectious diseases, we selected 1,331 individuals of European origin, as defined by principal component analysis (PCA) on WES data, as previously described (Belkadi, PNAS 2016). Based on a refined PCA on these 1,331 individuals, together with the 404 European 1000G individuals, we identified three distinct subpopulations (SI Appendix, Fig. S1): “Northern Europeans” (N), “Middle Europeans” (M) and “Southern Europeans” (S). For the real data analysis we used the craniosynostosis WES dataset reported in (20) (see *Supplemental Data*).

## RESULTS

### Simulation study

We first investigated the properties of our case-only test through simulations on real exome data from the 1000G populations and a subset of our in-house exome HGID database. We performed analyses under the null hypothesis of no digenic interactions, for which we assessed type I errors. We also worked under the alternative hypothesis of a digenic interaction, for which we assessed statistical power under genetic effects of different magnitudes. In these analyses, we compared the case-only approach to the corresponding case-control approach, for various population stratification (PS) scenarios.

### Type I error analyses

#### Case-only design

We first performed our case-only test on an ethnically homogeneous population based on the 214 IBS+TSI 1000G South-European samples. After the application of quality control filters (see *Supplemental Data*), 1,588 genes for which at least 15% of individuals carried rare variants were included in the analysis, resulting in 1,260,067 interaction tests. In tests of all possible pairs of genes, we observed a moderate inflation of type I error to 0.00147 for *α* = 0.1% (Table 2), and 0.0535 for *α* = 5% (Table S1). LD has been identified as a possible cause of type I error inflation in case-only tests (35). We therefore assessed the possible effect of LD, by restricting our analysis to pairs of genes physically separated by a minimal distance *δ* (measured in Mb). Empirical type I errors decreased with increases in *δ* from 0.1 to 2 Mb (Table S2), and a type I error of 0.00121 was obtained at a nominal value *α* of 0.1% when δ=2 Mb (Table 2). The distributions of *p* values for tests of pairs of genes with δ <2 Mb was strikingly inflated (SI Appendix, Fig. S2). In particular, the 204 *p* values <10^−10^ observed in the full analysis were all due to tests involving pairs of genes with δ <2 Mb. Type I errors did not improve significantly for δ >2 Mb (data not shown). Globally, these results show that LD accounted for the lowest *p* values in the case-only test. The refined investigation of statistically significant pairs of genes located close together (680 with *p* <0.05 among 4,082 pairs with δ <2 Mb in the IBS+TSI cohort) would require a case-control design. Even a small number of controls might help to reveal the true nature of the statistical signals for these pairs, through an analogous control-only approach, which would detect only LD. Even so, after simple LD correction based on removing the pairs of genes with δ <2 Mb, type I errors remained slightly above the corresponding upper limit of the confidence interval. No further improvement was obtained by adjusting our tests for the first three principal components, consistent with the fact that the IBS and the TSI populations are very close.

**Table 2.** Empirical type I errors at a nominal value of α = 0.1% for the case-only and case-control tests in the absence of population stratification.

| Design | Model |  |  |  |  |
| --- | --- | --- | --- | --- | --- |
| | $Pg_0^a$ | $Pg_2^b$ | $Pg_2 + 3PC^c$ | $Pg_2 + C_{25}^d$ | $Pg_2 + C_{35}^e$ |
| Case-only<br>(IBS+TSI) | *0.00147*<br>[0.0009-0.00110] | *0.00121*<br>[0.0009-0.00110] | *0.00133*<br>[0.0009-0.00110] | 0.00109<br>[0.0009-0.00113] | 0.00108<br>[0.0009-0.00114] |
| Case-control<br>(IBS+TSI+GBR+FIN) | *0.00128*<br>[0.0009-0.00110] | *0.00128*<br>[0.0009-0.00110] | *0.00130*<br>[0.0009-0.00110] | 0.00107<br>[0.0009-0.00113] | 0.00103<br>[0.0009-0.00114] |

Note: Boundaries of the 95% confidence intervals are shown in brackets. Type I error values lying outside the 95% confidence interval's boundaries are in italic.
<sup>a</sup> All pairs of genes with >15% of carriers of variants with MAF<5%.
<sup>b</sup> Pairs of genes as $Pg_0$ but with genes apart by at least 2 Mb.
<sup>c</sup> Pairs of genes as $Pg_2$ with adjustment on the first three principal components.
<sup>d</sup> Pairs of genes as $Pg_2$ with >25% of carriers of variants with MAF<10%.
<sup>e</sup> Pairs of genes as $Pg_2$ with >35% of carriers of variants with MAF<15%.

#### Case-control design

We conducted an analogous investigation with a case-control design on an enlarged European population consisting of the 404 IBS+TSI+GBR+FIN 1000G samples, in order to have ∼200 cases and ∼200 controls. We first applied it in a population balanced scenario (Table 2), in which 1,563 genes were retained after the application of quality control filters (*see Supplemental Data*). No inflation due to LD (as expected in a case-control design) or PS (as expected for a balanced scenario) was observed. Nevertheless, the empirical type I error of 0.00128 at *α* = 0.1% indicated that slight inflation, similar to that observed for the case-only test, also occurred with this test (Table 2). Similar trends were observed at *α* = 5% (Table S1). We hypothesized that this inflation might be at least partly due to the small sample sizes in the contingency cells of Table 1. We tested this hypothesis by repeating the analyses for both the case-only and the case-control tests with more common variants and a larger number of carriers at the gene level (i.e., variants with a MAF < 10% and genes with carriage rates of at least 25%, and variants with a MAF < 15% and genes with carriage rates of at least 35%; Table 2). The type I error was clearly lower, and improved as the frequency of variants increased. For both tests, empirical type I errors were within the boundaries of the confidence interval for *α* = 0.1%, but remained slightly above the upper limit of this interval for *α* = 5% (Table S1).

#### Sample size investigation

We investigated the impact of contingency cell sample sizes and the number of tests on the case-only approach, by extending the previous scenario to two new settings with less stringent MAF thresholds. First, we conducted a case-only test for all genes carried by at least 5% rather than 15% of individuals in the IBS+TSI population. This strategy increased the number of genes retained to 5,563, and, after the removal of genes in LD, we tested a total of 15,465,141 pairs of genes and generated the QQ-plot for SI Appendix, Fig. S3. The type I error was moderately inflated (0.057) for *α* = 5% and there was a slightly conservative type I error value (0.00085) for *α* = 0.1% (Table S3). Finally, we simulated the data for one gene considered “rare” (at least 1% carriers, total of 11,470 genes) and another considered “common” (at least 15% carriers, total of 1,588 genes). Under this scenario, 16,951,106 pairs of genes were tested, and the QQ-plot for SI Appendix, Fig. S4 was generated. The type I errors of 0.053 and 0.00097 obtained were closer to the expected values of 5% and 0.1%, respectively (Table S3). These results suggest that the case-only test is reliable for investigating a large range of carrier frequencies provided that LD is taken into account.

#### Population stratification

We then investigated the effect of PS, again focusing only on genes for which at least 15% of the individuals in the study population were carriers and which were separated by at least 2 Mb. For the case-only test, we used the 212 IBS+CHS samples, and we assessed 1,248 genes, in 776,879 tests (see *Supplemental Data*). Type I errors were highly inflated (0.0143 for *α* = 0.1% and 0.1264 for *α* = 5%) (Table 3 and Table S4). The application of PS correction (adjustment for the first three principal components) brought empirical type I errors back down to levels very similar to those previously observed (0.0013 for *α* = 0.1% and 0.0550 for *α* = 5%). For the case-control test, we used the 412 IBS+TSI+CHS+CDX samples under an unbalanced population scenario, with 1,173 genes (see *Supplemental Data*). Inflated type I errors were also observed (0.0026 for *α* = 0.1% and 0.0687 for *α* = 5%), although the inflation less striking. Adjustment for principal components (0.0013 for *α* = 0.1% and 0.0548 for *α* = 5%) resulted in values similar to those for a situation without PS (Table 3 and Table S4). Thus, provided that the search space was limited to pairs of genes far enough apart to avoid LD and adjustment for PCs was applied when required, our case-only test yielded reasonable type I error rates, similar to those for the analogous case-control approach.

**Table 3.** Empirical type I errors at a nominal value of α = 0.1% for the case-only and case-control tests in the presence of population stratification.

| Design | PC adjustment |  |
| --- | --- | --- |
|  | No adjustment | 3PC |
| Case-only<sup>a</sup><br>(IBS+CHS) | *0.01432*<br>[0.0009-0.00113] | *0.00135*<br>[0.0009-0.00113] |
| Case-control<br>Balanced<br>(IBS+TSI+CHS+CDX) | *0.00132*<br>[0.0009-0.00113] | *0.00136*<br>[0.0009-0.00113] |
| Case-control<br>Unbalanced<br>(IBS+TSI+CHS+CDX) | *0.00257*<br>[0.0009-0.00113] | *0.00126*<br>[0.0009-0.00113] |

Note: Boundaries of the 95% confidence intervals are shown in brackets. Type I error values lying outside the 95% confidence interval's boundaries are in italic.
<sup>a</sup> Using pairs of genes with genes apart by at least 2 Mb.

### Power analyses

#### Average power scenario

Power studies were conducted on an enlarged European population consisting of 1,735 individuals from the four European 1000G populations (IBS, TSI, GBR, FIN) and 1,331 individuals from the in-house HGID database (see *Supplemental Data*). We first estimated an “average” power by testing all possible pairs of genes (scheme A, Table 4), each with at least 15% carriers and separated by at least 2 Mb. In total, 370,530 tests were performed in 10 replicates (see *Supplemental Data)*. Fig. 1 displays the results obtained for scenarios including one or no main genetic effect, corresponding to the most pertinent situations in which to search for a gene x gene interaction. Adjusted and non-adjusted curves were superimposed, indicating that this analysis, in a European population, was not affected by PS. In all situations, power was always greater for the case-only test than for the case-control test. For example, a power of 65% at *α* = 0.1% was obtained when *OR*_*I*_ = 5 and no main effects were considered, whereas a power of only 40% was obtained for the corresponding case-control test in the same conditions. Similar trends were observed when one main effect was present (Fig. 1 and SI Appendix, Fig. S5) and for assessments of power at *α* = 5% (data not shown).

**Table 4.** Description of the schemes used in the *Power* section of the *Results.*

|  | Schemes |  |  |  |
| --- | --- | --- | --- | --- |
|  | *A* | 2G | 2GS | 2GR |
| Genes tested | Genome-wide | 2 genes |  |  |
| Genes characteristics | All genes | Both common and non-stratified by population | Both common and stratified by population | One common and one rare non-stratified by population |
| $OR_j^a$ | {1,2} | {1,2} | {1,2} | {1,2} |
| $OR_k^a$ | {1,2} | {1,2} | {1,2} | {1,2} |
| $OR_I^b$ | {1, ..., 5} | {1, ..., 5} | {1, ..., 5} | {1, ..., 10} |

<sup>a</sup> $OR_j$ and $OR_k$ are the odds ratios for the main effect of the first and the second gene of each pair, respectively.
<sup>b</sup> $OR_I$ is the odds ratio for the interaction term of Eq. 1.

**Fig. 1.**
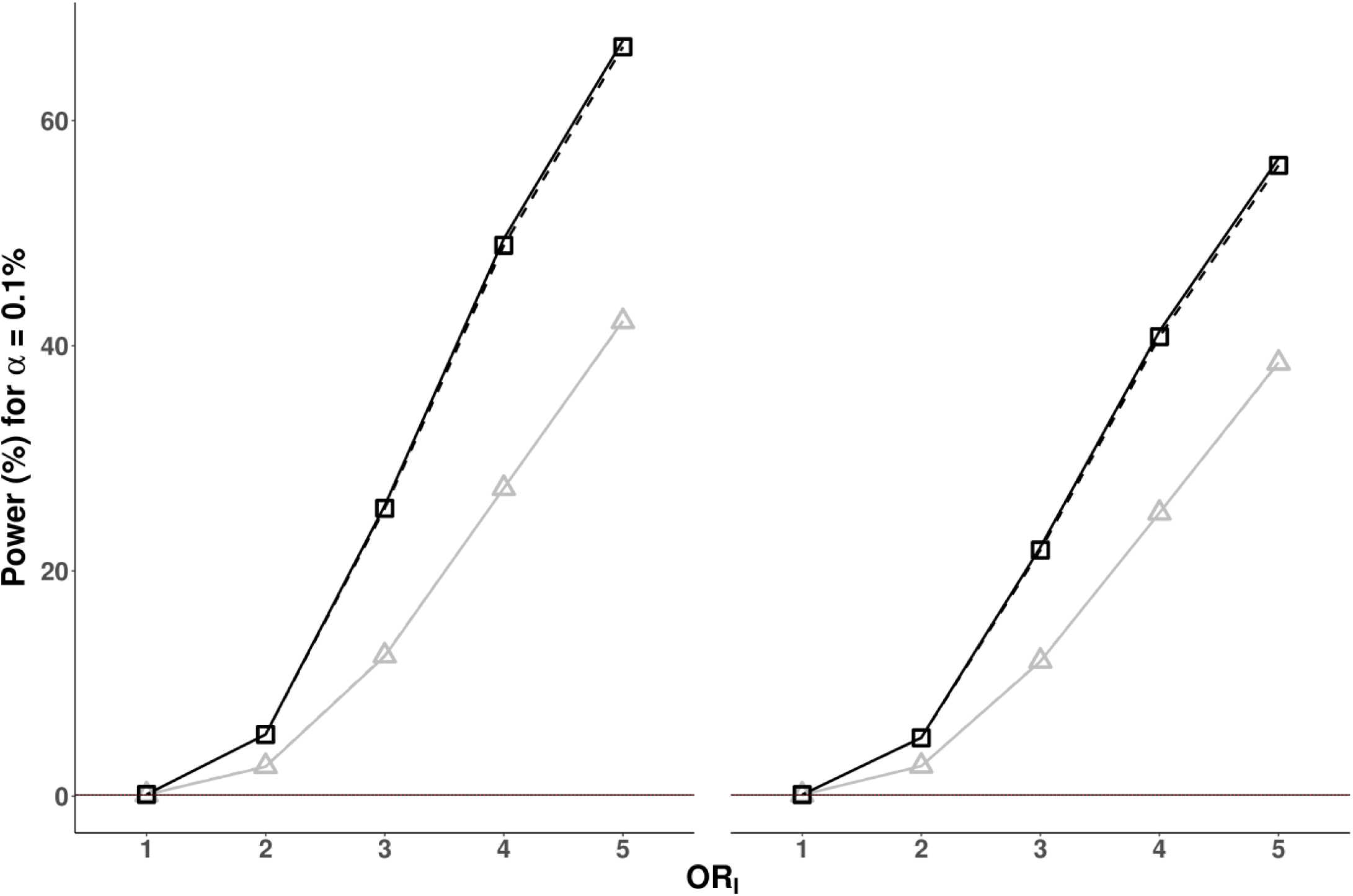
Power of the case-only and case-control tests for the analysis of all pairs of genes (scheme A). Power values are presented as a % for a type I error of 0.1%, as a function of the odds ratio for interaction (OR_I_), for the case-only (dark curves) and case-control (light curves) tests with (dotted lines with symbols) or without (solid lines without symbols) adjustment for the first three principal components. The left panel is obtained when no main gene effects are present, whereas the right panel shows results with a main effect of the second gene (OR=2). Note that the results with and without adjustment are very similar and the strong superimposition of the corresponding curves.

#### Two-gene power scenarios

We then focused on two specific pairs of genes, without (*AHNAK, PKHD1L1*, scheme 2G, see Table 4) and with (*ARPP21, MACF1*, scheme 2GS, see Table 4) PS (see *Supplemental Data*) In the analysis of scheme 2G, the case-only test performed better, overall, in terms of power (Fig. 2 and SI Appendix, Fig. S6, top figures). In the absence of main effects, with *OR*_*I*_ = 3 and α = 0.1%, a power value of 62% was obtained for the case-only test, versus only 27% for the case-control test.

**Fig. 2.**
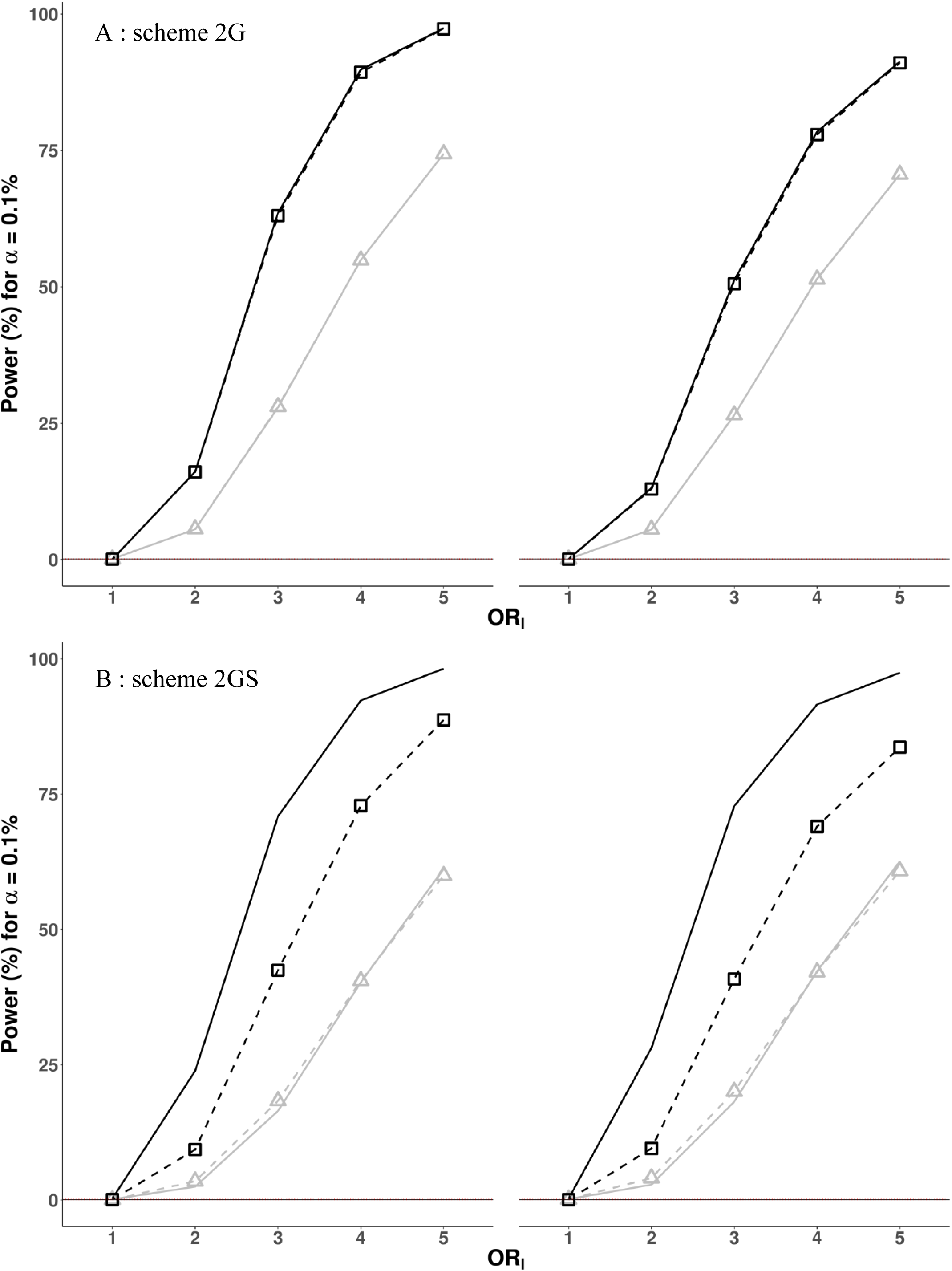
Power of the case-only and case-control tests for the analysis of two specific pairs of genes in the absence (scheme 2G) or presence (scheme 2GS) of population stratification. Power values are presented as in Figure 1. Results are shown for the analysis of A) the two non-stratified genes *PKHD1L1* and *AHNAK* (scheme 2G, top figure), and B) the two stratified genes *ARPP21* and *MACF1* (scheme 2GS, bottom figure). The left panel is obtained when no main gene effects are present whereas the right panel shows results with a main effect (OR=2) of the second gene, i.e. *AHNAK* and *MACF1* respectively.

For scheme 2GS, the power curves for the adjusted and non-adjusted case-only tests were not superimposed, indicating an effect of PS (Fig. 2 and SI Appendix, Fig. S6, bottom figures). We therefore used only the adjusted case-only test for comparison. As expected, the case-control test was not affected by PS (0.0009 for *α* = 0.1%) and had type I error values similar to those for the adjusted case-only test (0.0011 for *α* = 0.1%). The adjusted case-only test clearly outperformed the case-control test, by reaching a power of 90% when *OR*_*I*_ = 5 without main effects, for example, whereas the corresponding power for the case-control test was only 60%. Finally, we also considered another specific pair of genes, including one “common” (26% carriers) and one “rare” (5% carriers) gene (scheme 2GR, see Table 4). The case-only test was again more powerful than the corresponding case-control test (Fig. 3 and SI Appendix, Fig. S7), particularly in the absence of main effects, giving an absolute difference in power of almost 30% when *OR*_*I*_ = 10. Situations with a lower cumulative frequency of rare variants and a stronger OR might fit a Mendelian-like disorder hypothesis better and would be of particular interest concerning the application of this approach to real data presented below.

**Fig. 3.**
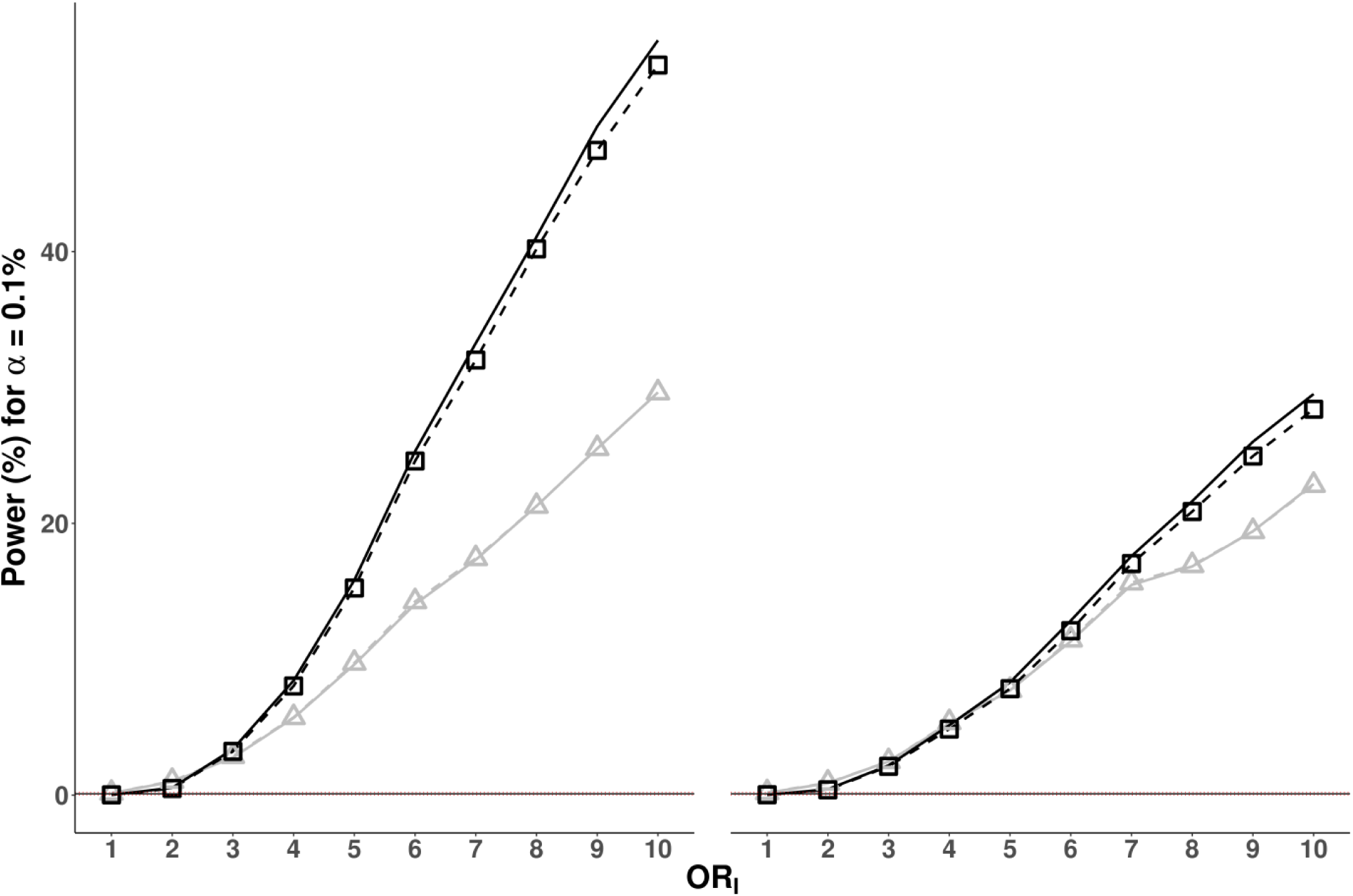
Power of the case-only and case-control tests for analyzing a pair of genes with different proportions of variant carriers (scheme 2GR). Power curves are presented as in Figure 1. Results are shown for the analysis of one “common” (*AHNAK*) and one “rare” gene (*MPC1*) (scheme 2GR). The left panel is obtained when no main effects are present, whereas the right pannel shows results with a main effect (OR=2) of the second gene, i.e. MPC1.

### Real data analysis: craniosynostosis

#### Background

We first applied our test to the dataset that led to the discovery of the first case of DI of non-syndromic midline craniosynostosis (MIM: 617439) (20). The original study showed a strong enrichment in rare heterozygous *SMAD6* mutations predicted to be damaging among cases (13 carriers among the 191 probands). Incomplete penetrance was observed in relatives of the carriers. The role of the common variant *rs1884302* (MAF=0.33 in European populations), located close to the *BMP2* gene and previously associated with craniosynostosis through GWAS (19), was therefore investigated, and this variant was found to account for almost all the observed phenotypic variation. Eleven of the 13 *SMAD6* probands were also carriers of *rs1884302*, whereas none of the healthy *SMAD6* carriers carried this variant. We used these data to determine whether our unbiased case-only test could detect this digenic association in the context of a GW search (i.e. without prior knowledge of the role of the *SMAD6* and *BMP2* variants).

#### Genome-wide search

In total, 285,216 tests (83 genes and 8,102 variants) were conducted on the WES data for 191 patients after the application of quality control and other filters to the variants and genes (see *Supplemental Data*). The resulting QQ-plot shows no deviation from the expected distribution, with only one significant result over the expected *p*-value line (Fig. 4). This result (*p* = 1.58×10^−6^, OR = 30.95) corresponds to the digenic combination of *SMAD6* and *rs1884302*, and is one order of magnitude higher than the second result (*p* = 1.04×10^−5^), which is close to the expected line. The 2×2 contingency table for the top result is shown in Table S5, and corresponds to the distribution found in the original paper (20). Thus, the two-locus genome-wide analysis focusing on genes harboring rare variants together with the potential contribution of a common modifier variant was able to detect the previously reported DI for craniosynostosis (20). This analysis provides proof-of-concept that our statistical test can detect DI without the need for biological assumptions concerning the disease studied, even when the disease is very rare.

**Fig. 4.**
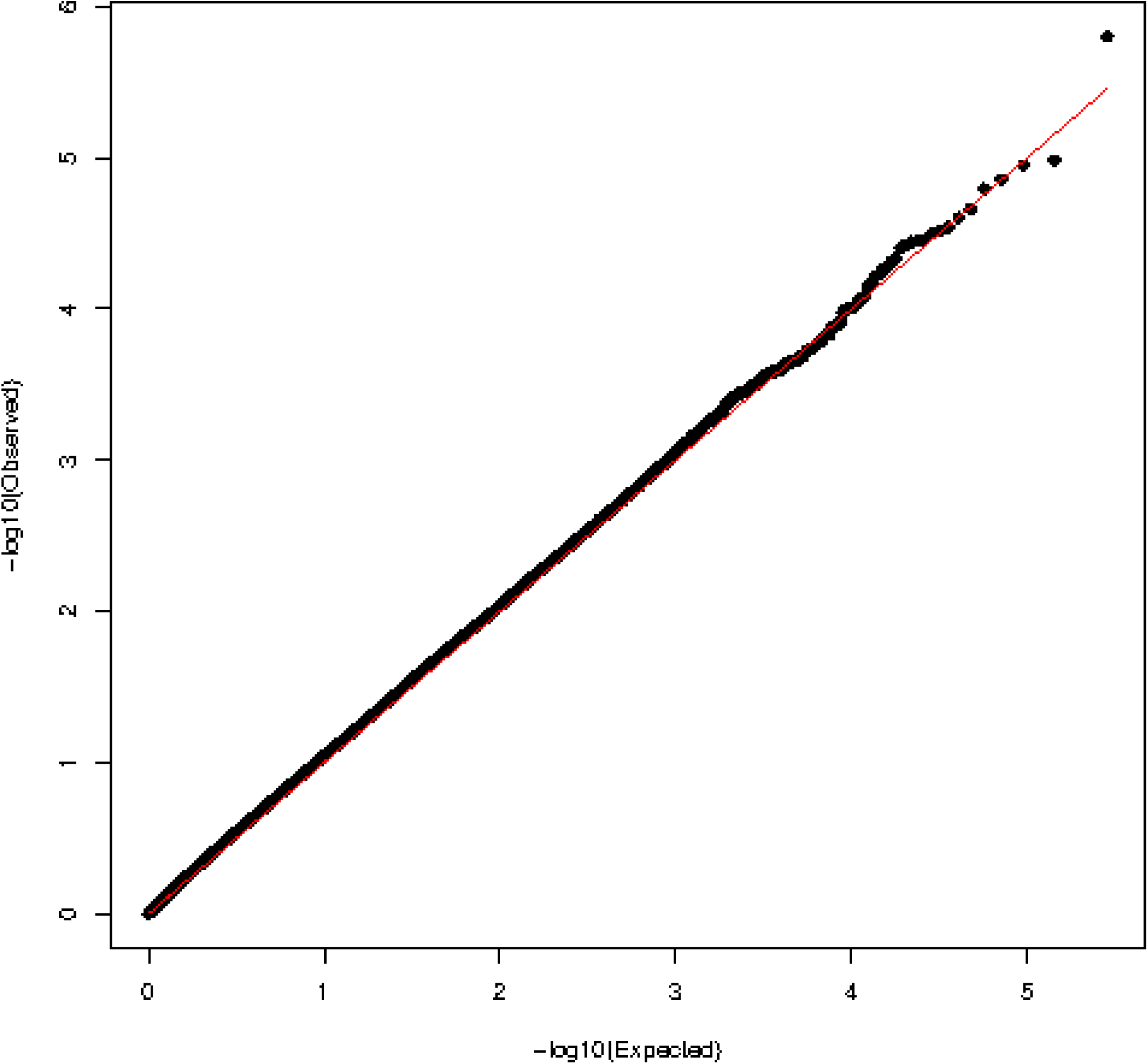
QQ-plot for the genome-wide case-only test conducted on the 191 craniosynostosis probands. QQ-plot for a genome-wide analysis under a dominant mode of inheritance, adjusted for the first three principal components, and considering pairs of genes and variants at least 2 Mb apart with > 5% carriers of rare variants a world-wide frequency > 10% for the variant (*n* = 285,216 pairs).

## DISCUSSION

There is increasing evidence to suggest that DI plays an important role in the genetic architecture of many conditions. The three previously reported approaches searching for gene x gene interactions in the general context of rare variant association studies are based on case-control designs (26–28). Moreover, these tests were assessed in limited simulation studies involving short genomic sequences of less than 500 variants (n=1) or only 20 variants (n=2), and were not based on WES-based simulated data. None was reported to have detected two genetic lesions at the GW level. Indeed, all previously successful DI studies relied on candidate gene approaches to overcome the lack of appropriate statistical resources to search for DI at the GW level (17). DI studies and statistical interaction approaches have thus been following separate paths. We show here, through both extensive simulation studies on real WES datasets and application to the example of craniosynostosis, that our method is robust and powerful for the identification of digenic lesions at the GW level. Our unbiased genetic confirmation of the reported digenic lesions in the craniosynostosis dataset composed only of exome data from cases, a common feature of real datasets for rare disorders, justifies the choice of a case-only test based on the aggregation of rare variants. Further strong support for this approach is provided by the higher overall power for the case-only approach than for the corresponding case-control test, as shown here, for the same number of cases. We present here the results for cohorts of at least 200 cases. We recommend using at least 100 cases to ensure sufficient statistical power, but this is not an absolute requirement as it depends on the proportion of double carriers among cases (strength of the genetic association).

The proposed methodology is simple to apply and flexible. It requires only the definition of a set of variants for testing, with filters based on features including MAF, variant annotations, and genetic models, defined before the analysis. It can, of course, be used at the gene level for the two loci studied. It can also directly assess the role of common variants as potential modifiers of a known monogenic defect. This assessment is achieved by simply replacing the gene by the variant as the unit of analysis, as illustrated in the craniosynostosis example. Our result also provide proof-of-concept that incomplete penetrance in disorders considered to be monogenic can be explained by a unique digenic combination. The frequency of carriers considered in our simulation studies may appear to be too high, but two important points must be taken into account when studying a rare disorder. First, these thresholds correspond to a cumulative frequency of the variants potentially contributing to the disease. The frequency of each individual allele may be much lower. Second, enrichment in the true disease-causing alleles would be expected in patients. For example, in the craniosynostosis dataset, the cumulative frequency of carriers of rare damaging *SMAD6* mutations is 6.8% (13 of 191), whereas the maximum frequency of carriers of these variants in gnomAD, which includes data from more than 50,000 individuals, is 0.01%. The proposed case-only test thus already appears to be a novel, powerful, and timely tool for detecting DI based on NGS data at the GW level in disorders that are not explained or only partly explained by a monogenic lesion.

## ACKNOWLEDGEMENTS

We would like to thank the patients and their families, whose cooperation was essential for collection of the data used in this study. We thank all members of the Laboratory of Human Genetics of Infectious Diseases for helpful discussions. Céline Desvallées, Tatiana Kochetkov, Dominick Papandrea, Cécile Patissier, Mark Woollett, Amy Gall and Yelena Nemirovskaya for their assistance.

## URLs

DIDA, http://dida.ibsquare.be/

OMIM, https://www.omim.org

gnomAD, https://gnomad.broadinstitute.org/

snpEff, http://snpeff.sourceforge.net/

