## Supplemental Data for "A genome-wide case-only test for the detection of digenic inheritance in human exomes"

**This PDF file includes:**

Supplementary text
Figures S1 to S7
Tables S1 to S7
SI References

**Supplementary Information Text**
**Supplemental methods**

### Simulation study

#### *Variant filtering and mode of inheritance*

WES was performed on our sample of 1,331 individuals with the exome capture SureSelect Human All Exon V4+UTRs. We focused on coding variants covered by this kit, with the following quality control criteria (1): depth of coverage (DP) > 8, genotype quality (GQ) > 20, minor read ratio (MRR) < 0.2, and call-rate > 95%. Data from samples of 1000G populations were left unchanged. Only diallelic variants from autosomal regions predicted by software SnpEff (2) to be missense or loss of function (LOF), and with the gnomAD (3) bioinformatics filter status of "PASS" were retained for the analysis. For rare variants, unless explicitly stated otherwise, we retained variants with a MAF<5% in the 1000G database. All analyses presented in this simulation followed a dominant mode of inheritance for each gene, i.e., carriers were defined as individuals harboring at least one copy of at least one allele within the unit of a gene.

#### *Type I error analyses*

*Case-only design*. Type I error investigations were performed with the six populations from the 1000G database. First, the case-only test was conducted by considering as affected all individuals from both the IBS+TSI (*n*=214) samples, presenting modest levels of PS. Affected individuals were then taken from both the IBS+CHS (*n*=212) samples, displaying strong levels of PS. We avoided sample size-related issues, by focusing on genes for which at least 15% of the study population was carrier of rare variants. Therefore, after the application of quality control filters (see *Variant filtering and mode of inheritance*), 1,588 and 1,248 genes were included in the

IBS+TSI and IBS+CHS analyses, resulting in 1,260,067 and 776,879 interaction tests, respectively.

*Case-control design*. The case-only test was compared with the described classic case-control test. We defined two combined 1000G samples, IBS+TSI+GBR+FIN (*n*=404) and IBS+TSI+CHS+CDX (*n*=412), composed of four European (EUROP sample) and two European and two Asian populations (EURAS sample), respectively. Half the individuals were simulated as cases or controls. We introduced a degree of PS, by considering balanced and unbalanced population scenarios for the definition of case-control status when analyzing the EURAS sample. In the balanced scenario, 50% of individuals in each of the four constitutive samples were considered, at random, to be cases or controls. In the unbalanced scenario, 5/6 of the individuals of the IBS+TSI samples were considered to be cases (and 1/6 controls), and 1/6 of the CHS+CDX individuals were simulated as cases (and 5/6 as controls). As we focused on genes with carriage rates of at least 15%, 1,563 genes were retained for the balanced case-control EUROP analysis, whereas 1,173 genes were used in the balanced and unbalanced EURAS analyses.

***Power analyses***

*Framework***.** For tests of power and to broaden our database for the independent sampling of cases, we used a final set of 1,735 European individuals combining the four European 1000G populations (IBS, TSI, GBR, FIN) with 1,331 individuals from the HGID database. For a given pair of genes, phenotype assignment followed a two-step procedure. First, the probability of being affected (penetrance) was calculated with the logistic regression model described in Eq 1. For a given predefined triplet $(\beta_j, \beta_k, \beta_I)$, accounting for main and interaction effects, the "baseline penetrance" parameter $\beta_0$ was computed to ensure 15% of cases, and examples of the resulting penetrances are shown in Table S6. Second, as individuals were classified according to their double-genotype status (four groups), phenotype was randomly assigned for a given individual based on the corresponding penetrance, giving a mean of 260 cases per replicate (15% of cases). We conducted simultaneous case-only and case-control tests, such that, for each replicate, the same number of controls as of simulated cases was sampled randomly from the control group.

*Average power scenario*. We first estimated an "average" power by testing all possible pairs of genes (scheme A, Table 4), focusing on genes carried by at least 15% of the sample of 1,735 individuals. With respect to type I error results, we only considered genes separated by at least 2 Mb, resulting in the analysis of a total of 253 genes (37,053 pairwise combinations). As there were 10 replicates, 370,530 tests were conducted. In addition, several models were considered according to the level of the interaction effect (OR from 1 to 5) and the presence or absence of main effects of either of the two genes (Table 4).

*Two-gene power scenarios*. We then focused on two specific pairs of genes, (*AHNAK, PKHD1L1*, scheme 2G, see Table 4) and (*ARPP21, MACF1*, scheme 2GS, see Table 4) with carriage rates of 26%, 33%, 17% and 36%, respectively and conducted 10,000 replications under the scenarios described above. These genes were chosen on the basis of their level of stratification across the three European subpopulations. As shown in Table S7, the first pair (*AHNAK, PKHD1L1*) of genes had similar frequencies across populations, whereas the second pair had different frequencies across populations

(*ARPP21, MACF1*). Accurate measurements were made by classical logistic regression analysis testing the correlation between PC1 (strongly associated with the north-south gradient of European ancestry, as shown in Figure S1) and the vector of carriers $G_j$ (Eq. 1) for each gene $j$ (Table S7). Finally, we considered a third pair of genes, *AHNAK* and *MPC1* (scheme 2GR, see Table 4), with a lower cumulative frequency of rare variants (5%) (Table S7). In this scheme, tests were performed under the settings described above, but with a larger range of interaction ORs (from 1 to 10 rather than 1 to 5, Table 4).

***Real data analysis: craniosynostosis***

Using the exome data for the 191 probands of the craniosynostosis dataset (4), we performed a digenic analysis assuming that rare variants at a first locus interacted with a common variant at a second locus. For this analysis, we considered only missense or predicted LOF variants, together with *rs1884302*, not captured by exome data. Indeed, the idea was to test whether a common variant leading to an effect like that observed in the craniosynostosis study could be detected in this digenic genome-wide analysis. We applied the same quality control filters to this exome set as described in the section on variant filtering, together with the blacklist strategy to remove additional non-relevant variants (5). We selected rare variants (world-wide gnomAD allele frequency $<10^{-4}$) for the first locus, and common variants (world-wide gnomAD allele frequency >0.1) for the second locus. For the first locus, only genes with at least 5% rare variant carriers were retained, to ensure sufficient power. We accounted for LD by using only pairs of variants and genes separated by at least 2 Mb for the analysis. For the second locus, we also pruned out variants in strong LD ($r^2$ >0.6), using sliding windows of 100 kb. We applied a dominant inheritance model for both loci, and *p* values were adjusted for the first three principal components.

**Fig. S1. Principal component analysis (PCA) on the European individuals used in the power simulation study**

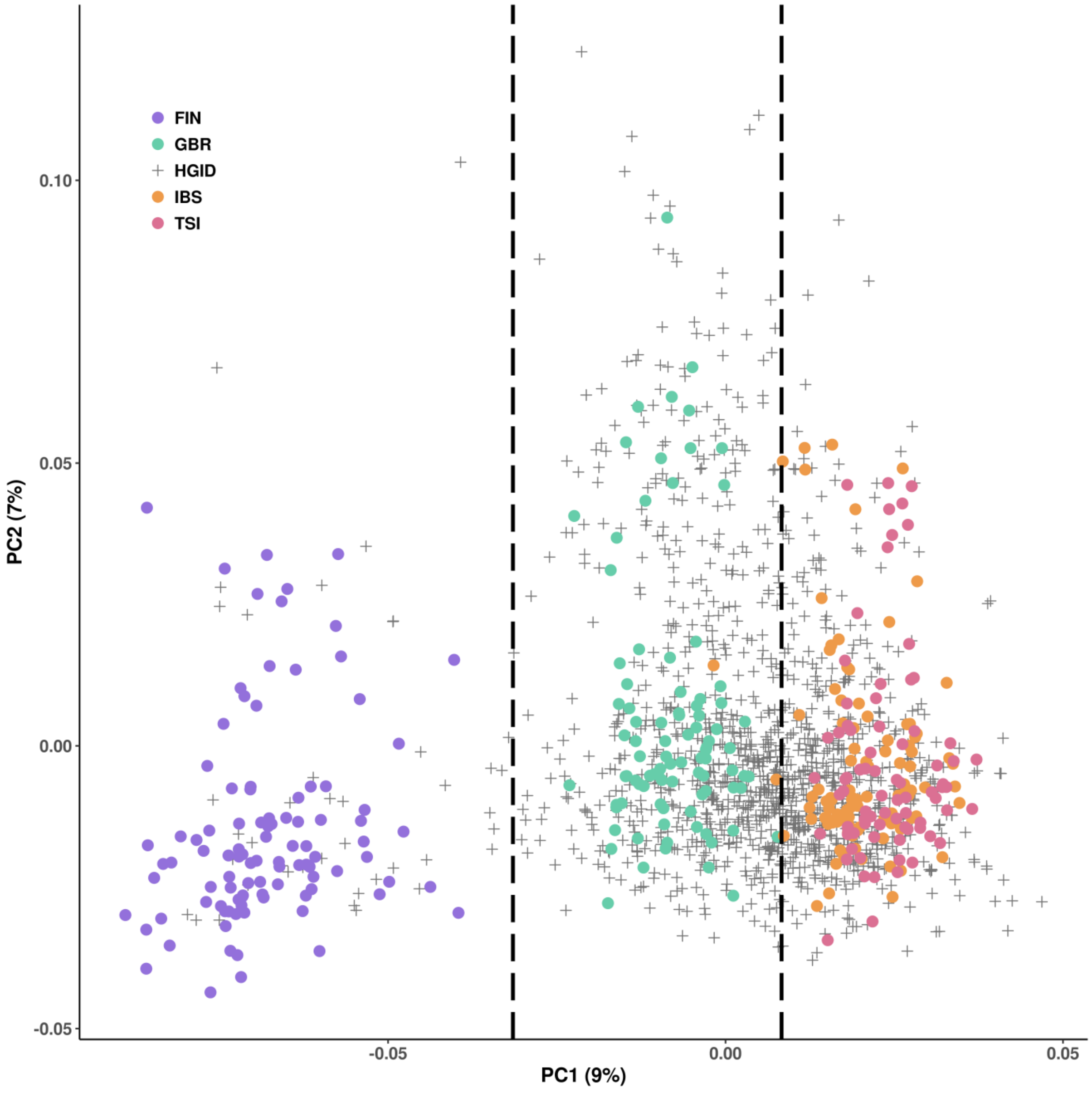


PCA (PC1 vs. PC2) conducted on WES data from 1,735 Europeans including 404 1000G subjects (FIN, GBR, IBS and TSI) and 1,331 subjects from our HGID database. Colors are used to represent the four subpopulations from the 1000G project (circles) and the individuals from the HGID cohort (gray crosses). Two vertical dotted lines were drawn along PC1 based on the classical north-south gradient of European ancestry, further defining three subpopulations: “Northern Europeans” (including FIN), “Middle Europeans” (including GBR) and “Southern Europeans” (including IBS and TSI).

**Fig. S2. QQ-plots for the case-only test conducted on the IBS+TSI population, as a function of distance between the two genes considered**

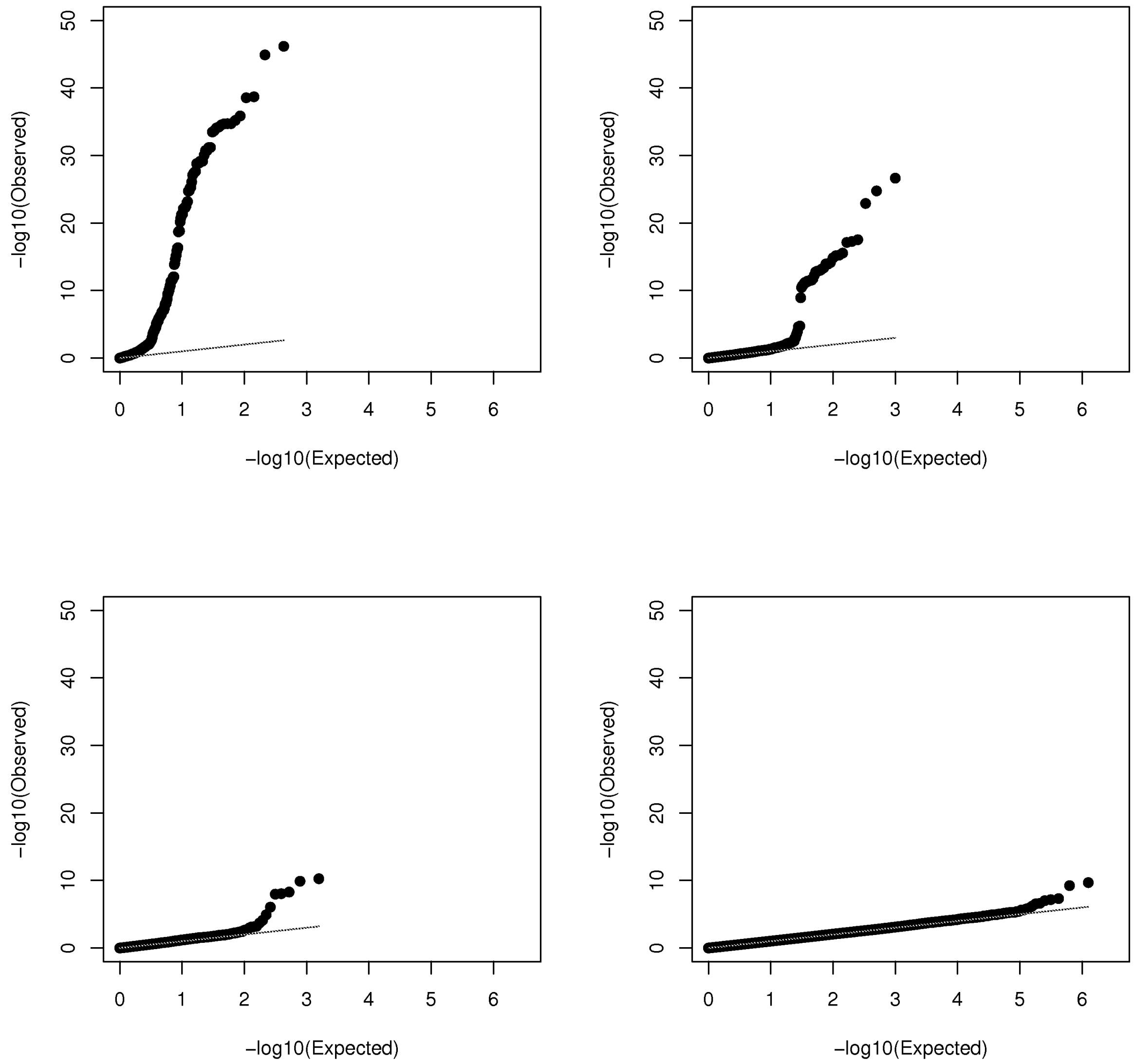


QQ-plots are shown for the case-only test on the IBS+TSI population, with distances between the two genes of a pair of (A) < 0.1 Mb, (B) between 0.5 and 1Mb, (C) between 1 and 2 Mb, and (D) > 2 Mb.

**Fig. S3. QQ-plot for the genome-wide case-only test conducted on the IBS+TSI population with a 5% variant carrier threshold for the first and second gene of each pair**

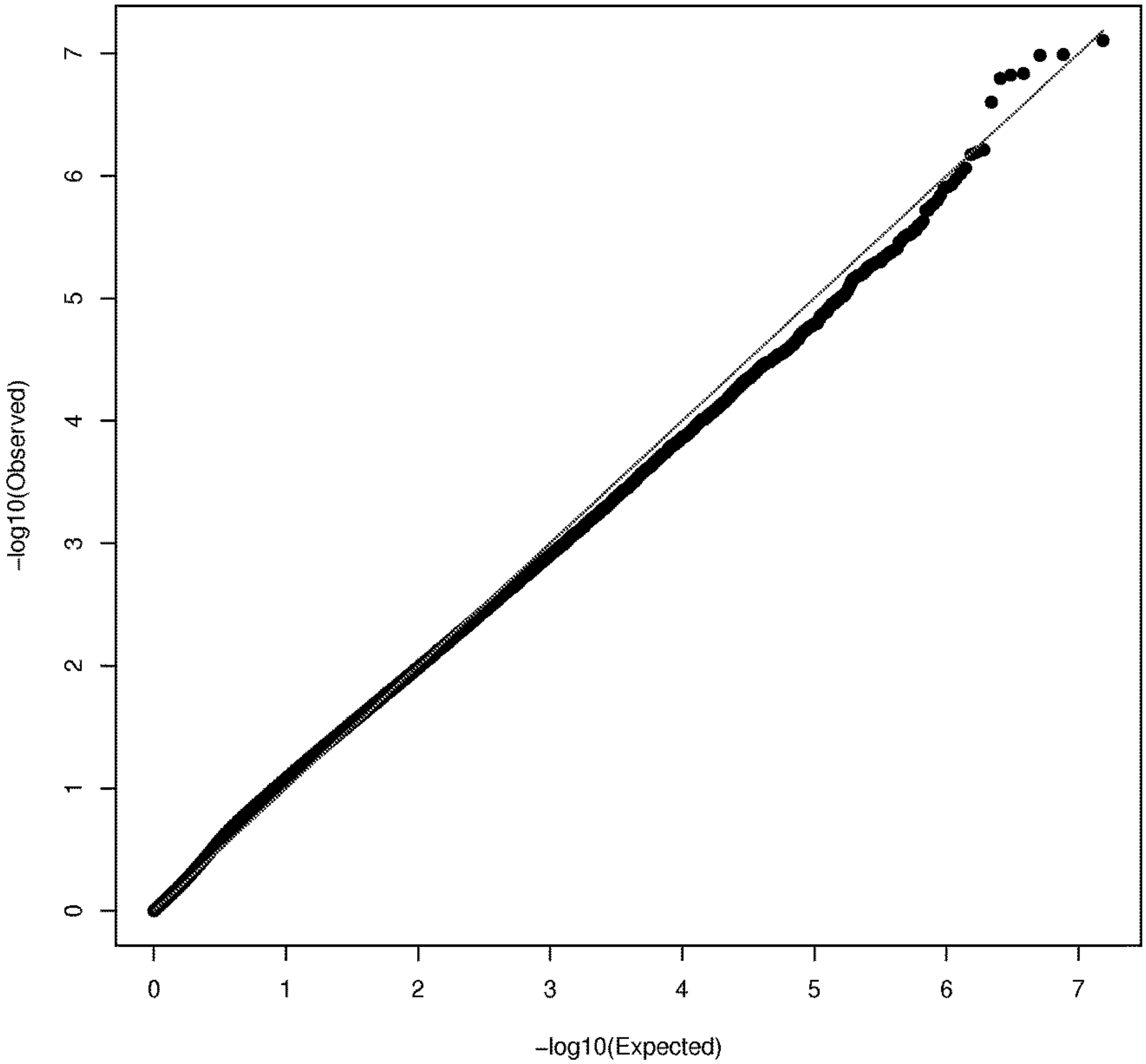


QQ-plot for pairs of genes more than 2 Mb apart.

**Fig. S4. QQ-plot for the genome-wide case-only test conducted on the IBS+TSI population with a variant carrier threshold of 1% and 15% for the first and second genes of each pair, respectively**

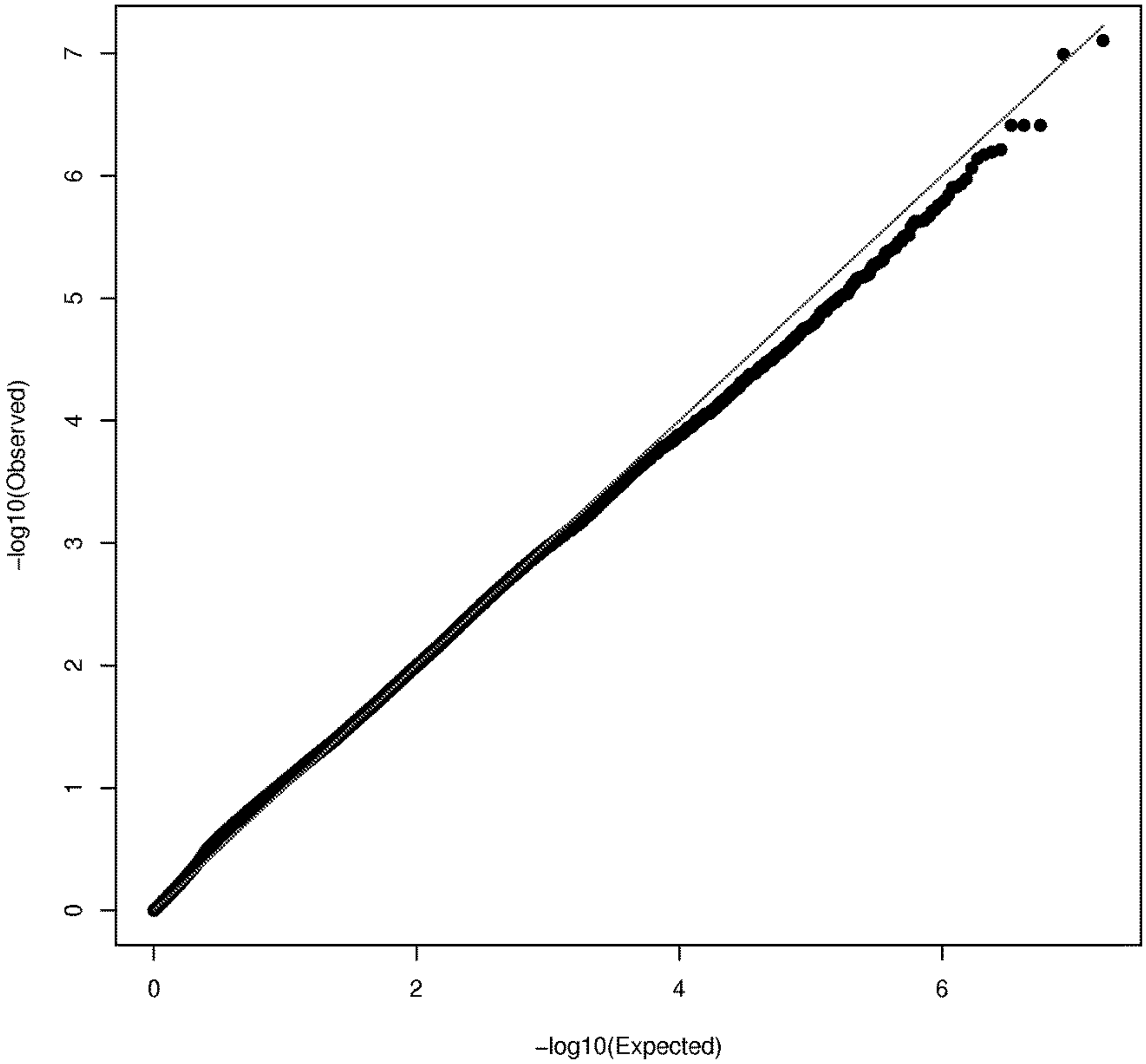


QQ-plot as in Figure S3.

**Fig. S5. Power of the case-only and case-control tests for analyzing all pairs of genes (scheme A)**

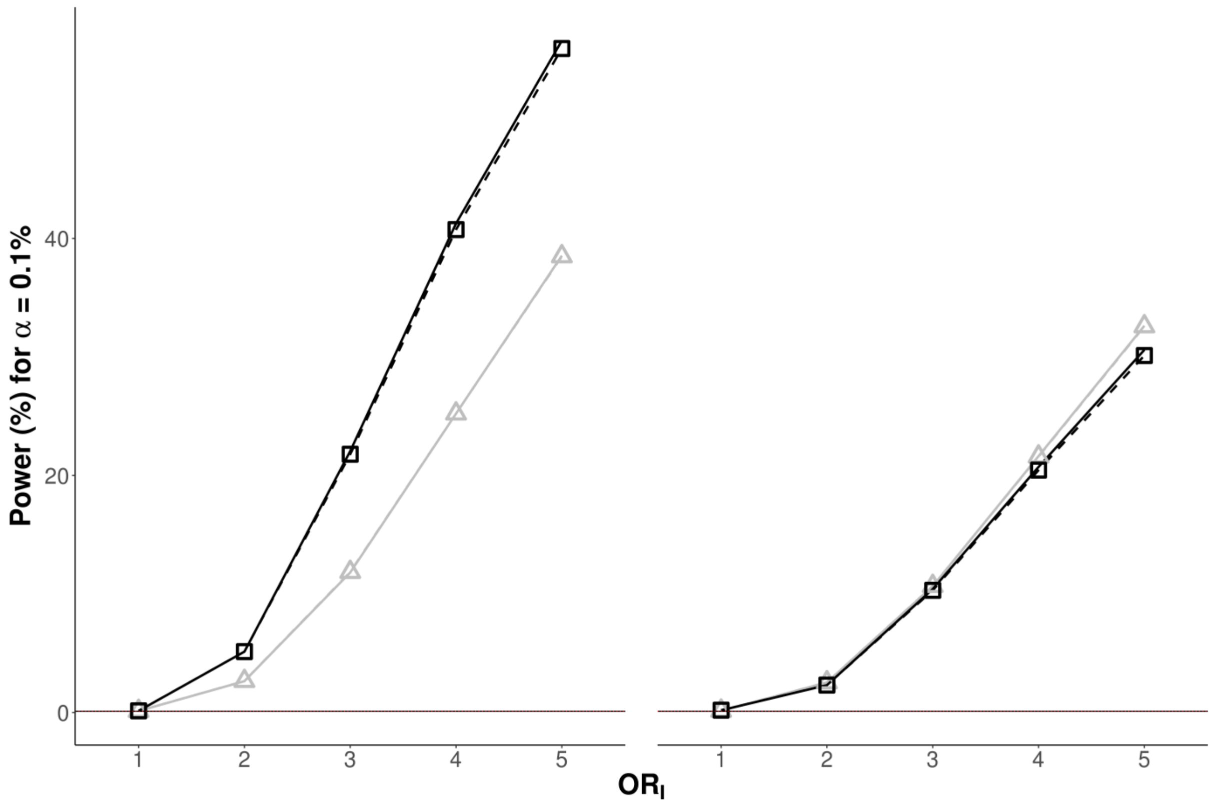


Power values are presented as a %, for a type I error of 0.1%, as a function of the odds ratio for interaction, ($OR_I$), for the case-only (dark curves) and case-control (light curves) tests with (dotted lines with symbols) and without (solid lines without symbols) adjustment for the first three principal components. The left panel is obtained when a main effect of the first gene is present (OR=2) whereas the right panel shows results for main effects of both genes (OR=2 for both genes).

**Fig. S6. Power of the case-only and case-control tests for analyzing two specific pairs of genes in the absence (scheme 2G) or presence (scheme 2GS) of population stratification**

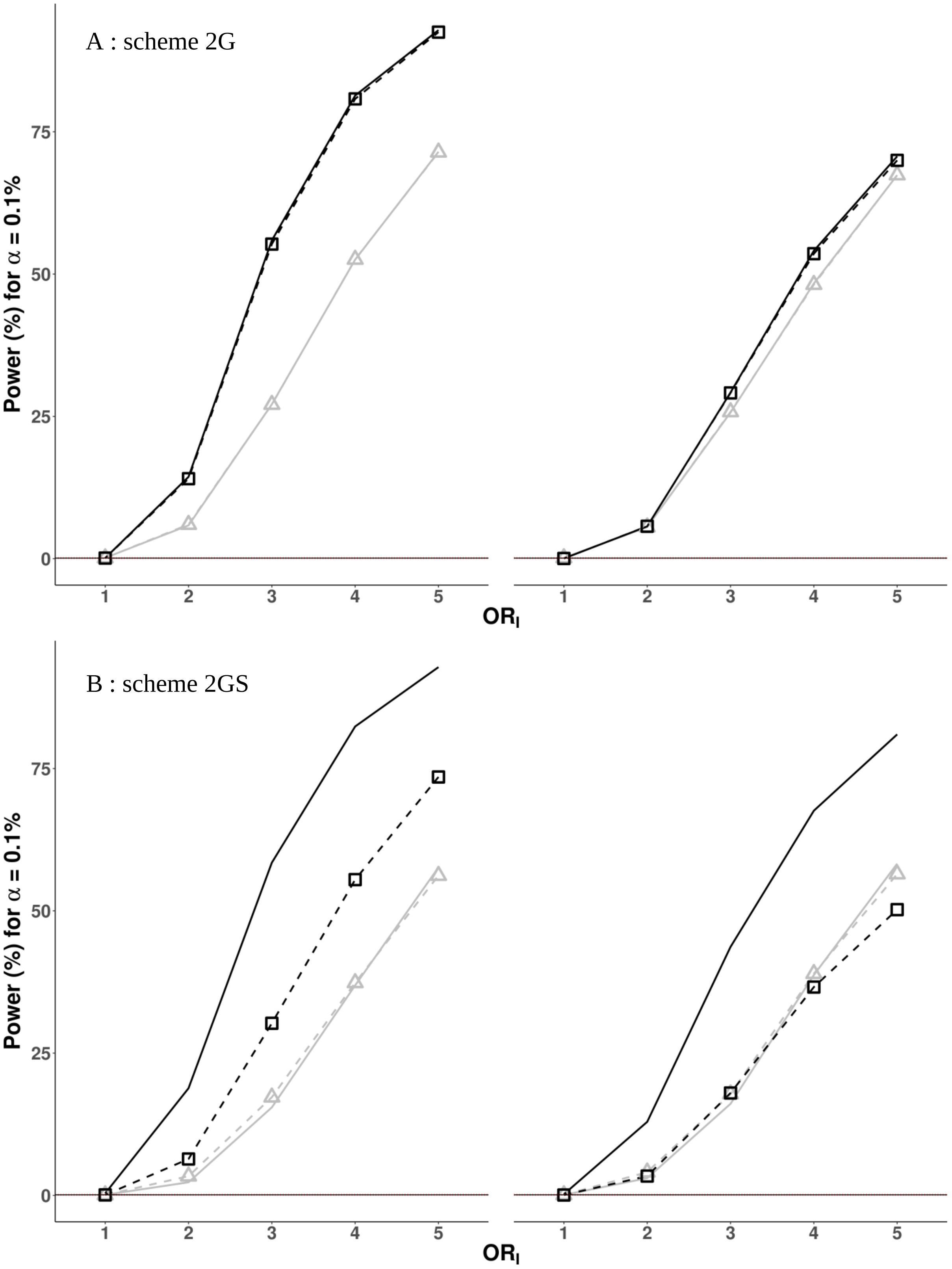
A : scheme 2G
B : scheme 2GS
Power (%) for α = 0.1%
0
25
50
75
1
2
3
4
5
OR$_I$

Power values are presented as in Figure S5. The left panel is obtained when a main effect of the first gene is present (OR=2; i.e. *PKHD1L1* and *ARPP21*), whereas the right panel shows results with main effects of both genes (OR=2 for both genes).

**Fig. S7. Power of the case-only and case-control tests for analyzing a pair of genes with different proportions of variant carriers (scheme 2GR)**

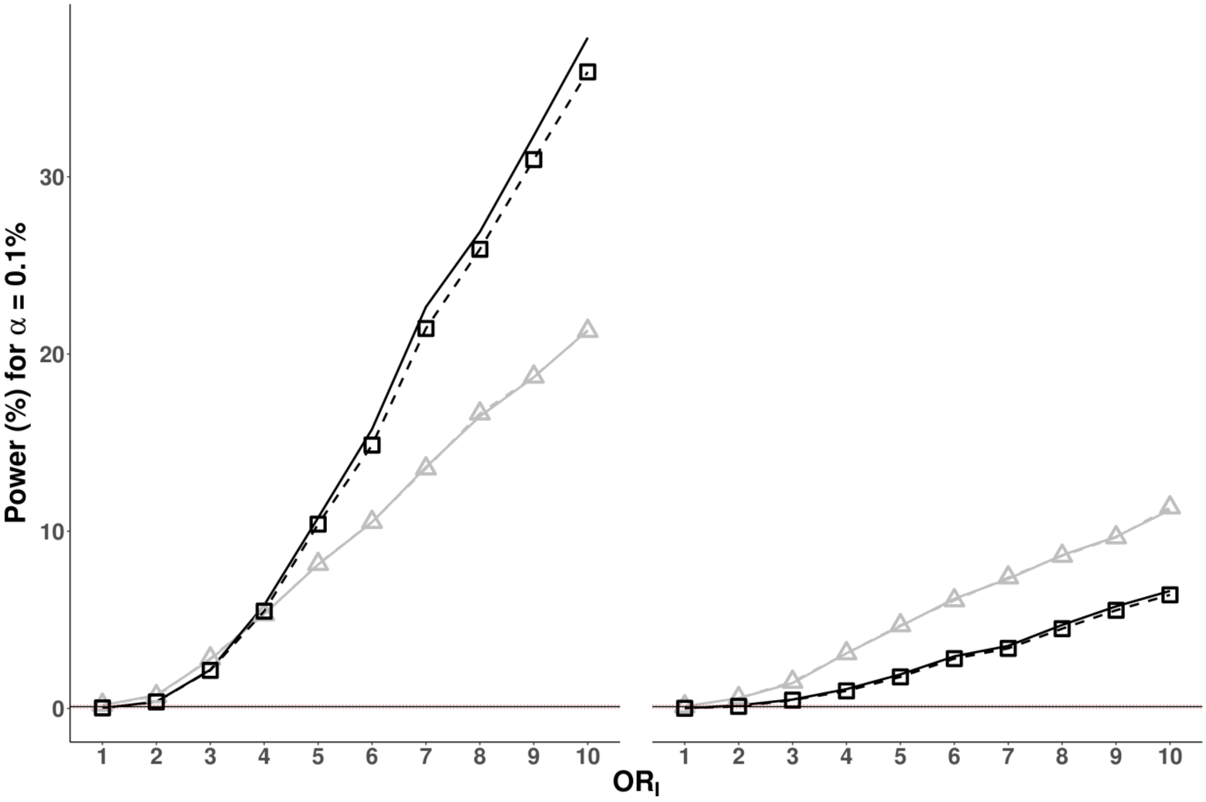


Power values are presented as in Figure S5. The left panel is obtained when a main effect of the first gene is present (OR=2; i.e. *AHNAK*), whereas the right panel shows results with main effects of both genes (OR=2 for both genes).

**Table S1. Empirical type I errors at a nominal value of α = 5% for the case-only and case-control tests in the absence of population stratification.**

| | Model | | | | |
|---|---|---|---|---|---|
| Design | $Pg_0$ [a] | $Pg_2$ [b] | $Pg_2 + 3PC$ [c] | $Pg_2 + C_{25}$ [d] | $Pg_2 + C_{35}$ [e] |
| Case-only (IBS+TSI) | *0.0535* [0.0494-0.0505] | *0.0531* [0.0494-0.0505] | *0.0549* [0.0494-0.0505] | 0.0512 [0.0494-0.0506] | 0.0511 [0.0494-0.0506] |
| Case-control (IBS+TSI+GBR+FIN) | *0.0541* [0.0494-0.0505] | *0.0541* [0.0494-0.0505] | *0.0544* [0.0494-0.0505] | 0.0516 [0.0494-0.0506] | 0.0518 [0.0494-0.0506] |

Note: Boundaries of the 95% confidence intervals are shown in brackets. Type I error values lying outside the 95% confidence interval's boundaries are in italic.

[a] All pairs of genes with >15% of carriers of variants with MAF<5%.
[b] Pairs of genes as $Pg_0$ but with genes apart by at least 2 Mb.
[c] Pairs of genes as $Pg_2$ with adjustment on the first three principal components.
[d] Pairs of genes as $Pg_2$ with >25% of carriers of variants with MAF<10%.
[e] Pairs of genes as $Pg_2$ with >35% of carriers of variants with MAF<15%.

**Table S2. Empirical type I errors at nominal values of α = 5% and α = 0.1% for the case-only test according to the minimal distance between the two genes of a pair.**

| | Case-only (IBS+TSI) Minimal distance (Mb) between genes of a pair | | | | |
|---|---|---|---|---|---|
| Type I error (%) | 0 | 0.1 | 0.5 | 1 | 2 |
| 5% | 0.05350 | 0.05337 | 0.05321 | 0.05317 | 0.05314 |
| 0.1% | 0.00147 | 0.00137 | 0.00126 | 0.00123 | 0.00122 |

**Table S3. Empirical type I errors at nominal values of α = 5% and α = 0.1% for the case-only test, for variant carrier thresholds other than 15% for the first and/or second gene of each pair.**

| | Carrier's threshold for gene 1 vs Carrier's threshold for gene 2 | |
|---|---|---|
| Type I error | 5% vs 5% | 1% vs 15% |
| α = 0.1% | *0.00085* [0.00100-0.00102] | *0.00097* [0.00100-0.00102] |
| α = 5% | *0.057* [0.0499-0.0501] | *0.053* [0.0499-0.0501] |

**Table S4. Empirical type I errors at a nominal value of α = 5% for the case-only and case-control tests in the presence of population stratification.**

| Design | PC adjustment | |
|---|---|---|
| | No adjustment | $3PC$ |
| Case-only[a] (IBS+CHS) | *0.1264* [0.0493-0.0506] | *0.0550* [0.0493-0.0506] |
| Case-control Balanced (IBS+TSI+CHS+CDX) | *0.0550* [0.0493-0.0506] | *0.0558* [0.0493-0.0506] |
| Case-control Unbalanced (IBS+TSI+CHS+CDX) | *0.0687* [0.0493-0.0506] | *0.0548* [0.0493-0.0506] |

Note: Boundaries of the 95% confidence intervals are shown in brackets. Type I error values lying outside the 95% confidence interval's boundaries are in italic.

[a] Using pairs of genes with genes apart by at least 2 Mb.

**Table S5. Contingency table for the digenic combination *rs1884302* (*BMP2*) – *SMAD6* in 191 craniosynostosis patients.**

| | | *SMAD6* | | |
|---|---|---|---|---|
| | | Carriers | Non carriers | Total |
| *BMP2* rs1884302 | Carriers | 11 | 38 | 49 |
| | Non carriers | 2 | 140 | 142 |
| Total | | 13 | 178 | 191 |

**Table S6. Penetrance values for the digenic combination of $G_1$ and $G_2$ when analyzing two specific pairs of genes with different (scheme 2GR) and similar (scheme 2G) proportions of variant carriers.**

| | Penetrances | | | | | | | |
|---|---|---|---|---|---|---|---|---|
| Scheme | $2G$ | | | | $2GR$ | | | |
| Carriers | $\overline{G_1} \times \overline{G_2}$[c] | $G_1$ | $G_2$ | $G_1 \times G_2$ | $\overline{G_1} \times \overline{G_2}$ | $G_1$ | $G_2$ | $G_1 \times G_2$ |
| $\boldsymbol{OR_2}^{\boldsymbol{a}} = \mathbf{1}$ | | | | | | | | |
| $\boldsymbol{OR_I}^{\boldsymbol{b}} = \mathbf{2}$ | 0.14 | 0.14 | 0.14 | 0.25 | 0.15 | 0.15 | 0.15 | 0.26 |
| $\boldsymbol{OR_I} = \mathbf{3}$ | 0.13 | 0.13 | 0.13 | 0.32 | 0.15 | 0.15 | 0.15 | 0.34 |
| $\boldsymbol{OR_I} = \mathbf{5}$ | 0.12 | 0.12 | 0.12 | 0.41 | 0.15 | 0.15 | 0.15 | 0.46 |
| $\boldsymbol{OR_I} = \mathbf{10}$ | - | - | - | - | 0.14 | 0.14 | 0.14 | 0.63 |
| $\boldsymbol{OR_2} = \mathbf{2}$ | | | | | | | | |
| $\boldsymbol{OR_I} = \mathbf{2}$ | 0.11 | 0.11 | 0.20 | 0.33 | 0.14 | 0.14 | 0.25 | 0.40 |
| $\boldsymbol{OR_I} = \mathbf{3}$ | 0.10 | 0.10 | 0.19 | 0.41 | 0.14 | 0.14 | 0.25 | 0.50 |
| $\boldsymbol{OR_I} = \mathbf{5}$ | 0.09 | 0.09 | 0.17 | 0.51 | 0.14 | 0.14 | 0.25 | 0.62 |
| $\boldsymbol{OR_I} = \mathbf{10}$ | - | - | - | | 0.14 | 0.14 | 0.24 | 0.76 |

Note: No main effect on the first gene ($OR_1 = 1$).

[a] $OR_2$ is the odds ratios for the main effect of the second gene of each pair.

[b] $OR_I$ is the odds ratio for the interaction term of Eq. 1.

[c] Individuals not carrying any rare variant in the first nor in the second gene.

**Table S7. Proportion (as a %) of variant carriers for the genes tested in the *Power* section of the *Results,* for the European population and subpopulations, as defined in *Material and Methods.***

| | | Carriers of European populations (%) | | | |
|---|---|---|---|---|---|
| P-value[a] | Gene | Global | Southern | Middle | Northern |
| 3.25E-32 | *ARPP21* | 17 | 12 | 16 | 45 |
| 2.05-05 | *MACF1* | 36 | 33 | 37 | 51 |
| 0.0049 | *MPC1* | 5 | 6 | 5 | 1 |
| 0.8937 | *AHNAK* | 26 | 26 | 27 | 23 |
| 0.9551 | *PKHD1L1* | 33 | 32 | 34 | 29 |

[a] Test for the homogeneity of the frequency of carriers across the three European subpopulations (see *Material and Methods*).
